# Alpha-band oscillations track the retrieval of precise spatial representations from long-term memory

**DOI:** 10.1101/207860

**Authors:** David W. Sutterer, Joshua J. Foster, John T. Serences, Edward K. Vogel, Edward Awh

**Author notes:** **Corresponding Authors:** David Sutterer or Edward Awh Institute for Mind and Biology 940 E 57^th^ St Chicago, IL 60637.

## Abstract

A hallmark of episodic memory is the phenomenon of mentally re-experiencing the details of past events, and a well-established concept is that the neuronal activity that mediates encoding is reinstated at retrieval. Evidence for reinstatement has come from multiple modalities, including functional Magnetic Resonance Imaging (fMRI) and electroencephalography (EEG). These EEG studies have shed light on the time-course of reinstatement, but have been limited to distinguishing between a few categories and/or limited measures of memory strength. The goal of this work was to investigate whether recently developed experimental and technical approaches, namely an inverted encoding model applied to alpha oscillatory power in conjunction with sensitive tests of memory retrieval in a continuous space, can track and reconstruct memory retrieval of specific spatial locations. In Experiment 1, we establish that an inverted encoding model applied to multivariate alpha topography can track retrieval of precise spatial memories. In Experiment 2, we demonstrate that the pattern of multivariate alpha activity at study is similar to the pattern observed during retrieval. Finally, we observe that these encoding models predict memory retrieval behavior, including the accuracy and latency of recall. These findings highlight the broad potential for using encoding models to characterize long-term memory retrieval.

## Introduction

Episodic memory is defined by the phenomenon of re-experiencing the details of past events, and is thought to be supported by the reactivation of neural activity that was present at encoding. In line with this view, functional magnetic resonance imaging (fMRI) studies have shown that sensory regions involved in the initial processing of information are re-engaged at retrieval (Wagner et al. 2005; Danker & Anderson, 2010), and that the voxel-wise patterns of activity within these regions resembles activity seen during encoding (Ritchey et al. 2013; Bosch et al. 2014; Hindy et al. 2016). In addition, more time-resolved measures of neural activity such as electroencephalography (EEG) and magnetoencephalography (MEG) have shown that retrieval-related neural activity echoes the broad strokes of encoding-related activity, such as the category of the paired associate (Wimber et al. 2012; Morton et al. 2013; Jafarpour et al. 2014; Waldhauser et al. 2016) or the task performed at encoding (Johnson et al. 2015). However, an open question is whether EEG activity can provide a temporally resolved means of tracking the retrieval of precise feature values that are associated with specific items.

Here, we addressed this question by measuring EEG activity during the encoding and recall of spatial information from long-term memory (LTM). To reconstruct the spatial representations in a precise manner, we applied an inverted encoding model (IEM) to the topography of oscillatory activity on the scalp. IEMs have provided a useful approach for reconstructing precise spatial representations from fMRI and EEG activity (Sprague and Serences 2013; Sprague et al. 2014, 2016; Foster et al. 2016; Foster, Sutterer, et al. 2017). However, this approach has not yet been applied to the study of long-term memory; therefore, it is an open question whether it is possible to track retrieval of spatial long-term memories by applying an IEM to EEG activity. Furthermore, it is unclear which frequency bands would carry this spatially specific information. On the one hand, previous work using an IEM applied to alpha-band EEG activity, has successfully tracked covert spatial attention (Foster, Sutterer, et al. 2017), and spatial representations maintained in working memory(Foster et al. 2016; Foster, Bsales, et al. 2017). Recent theories about the role of rhythmic oscillations in memory maintain that same frequencies of oscillations coordinate specific cognitive operations at encoding and retrieval (Siegel et al. 2012; Watrous and Ekstrom 2014; Watrous et al. 2015), predicting that alpha-band activity may play a similar role at retrieval. In addition, alpha-band activity has been shown to track hemifield-specific location memory (Stokes et al. 2012; Waldhauser et al. 2016). On the other hand, other frequency bands, especially theta and beta, are known to play important roles in long-term memory encoding and retrieval (Nyhus and Curran 2010; Morton et al. 2013; Hsieh and Ranganath 2014; Morton and Polyn 2017) and spatial navigation (Watrous et al. 2011; Bohbot et al. 2017). It is also possible that a combination of these frequencies carry spatially specific information. Thus, we investigated whether precise spatially specific information is reinstated during long-term memory retrieval, and, if so, which frequency bands carry this information.

In two experiments, participants learned to associate objects with specific angular locations. Then, they were asked to precisely report the associated location when presented with an object cue. This continuous recall procedure enabled a fine-grained measurement of mnemonic performance, and recent work has shown that modeling of the response error distribution can provide robust indices of the probability and precision of the stored representations (Zhang and Luck 2008; Harlow and Yonelinas 2014; Richter et al. 2016; Sutterer and Awh 2016). The IEM analysis revealed that spatially specific oscillatory activity tracked the retrieved locations after the presentation of the object cue. Consistent with oscillatory reinstatement accounts, we primarily observed spatially specific patterns of activity in the alpha band, just as observed in past studies of spatial working memory (Foster et al. 2016; Foster, Bsales, et al. 2017). Moreover, the alpha-band patterns observed during retrieval matched those observed during the initial encoding of the objects, in line with the hypothesis that encoding-related oscillatory patterns were reinstated during retrieval from LTM. Finally, the selectivity of alpha-band activity tracked memory performance as learning progressed as well as the latency with which participants reported the target locations. Together these findings suggest that LTM retrieval yields a reinstatement of the spatially specific oscillatory activity that is observed during encoding, and that multivariate analysis of alpha-band activity provides a powerful measure of the timing and success of this basic cognitive process.

## Materials and Methods

**Participants.** Sixty-nine adults (33 in Experiment 1, and 36 in Experiment 2; 18–35 years old, 38 female) participated in the study for monetary compensation ($10 per hour in Experiment 1, and $15 per hour in Experiment 2). All participants reported normal or corrected-to-normal vision and provided informed consent according to procedures approved by the University of Oregon Institutional Review Board (Experiment 1) and the University of Chicago Institutional Review Board (Experiment 2).

**Participant exclusions for Experiment 1.** For Experiment 1 participants were excluded for poor performance on the task and excessive EEG artifacts. One participant did not return for the second day of the experiment. One participant was excluded for poor performance on the first day (86.1° average response error across all day 1 tests) and data collection was terminated for one participant during the session for excessive artifacts. In addition, participants were excluded from further analysis if they had insufficient artifact-free trials (<550 trials). Artifact number exclusion criteria were set during data collection, but before the data were analyzed. Three participants were excluded due to excessive EEG artifacts. In the final sample, there were 27 participants in Experiment 1 (mean number of artifact-free trials = 799, *SD* = 85).

**Participant exclusions for Experiment 2.** For Experiment 2, our target final sample size was 24 subjects. Participants were replaced for poor task performance or if too many trials were lost due to recording or ocular artifacts. One participant was excluded for poor performance on LTM trials (87.1° average response error across all retrieval tests), and data collection was terminated for three participants during the session for excessive artifacts. In addition, participants were excluded from further analysis if they had insufficient artifact-free trials (<450 trials for encoding or retrieval). Artifact number exclusion criteria were set during data collection, but before the data were analyzed. We relaxed the exclusion criterion in Experiment 2 because we obtained fewer trials per condition. Eight participants were excluded due to excessive EEG artifacts. In the final sample there were 24 participants in Experiment 2 (mean number of artifact-free encoding trials = 535, *SD* = 46 and recall trials = 545, *SD* = 39).

**Apparatus.** Stimuli were presented in MATLAB using Psychtoolbox (Brainard, 1997; Pelli, 1997) and were presented on a 17-in. CRT monitor (60 Hz) for Experiment 1 and on a 24-in. LCD monitor (120 Hz) for Experiment 2.

**Experiment 1 task procedure.** The experiment was comprised of two sessions run on consecutive days (Figure 1a). On Day 1, participants were instructed to learn 120 object-location associations (see Figure 1a for example clip art) as accurately as possible for the test the next day. On Day 2, participants were cued with the object and asked to recall and report the associated location while we recorded EEG data.

**Figure 1.**
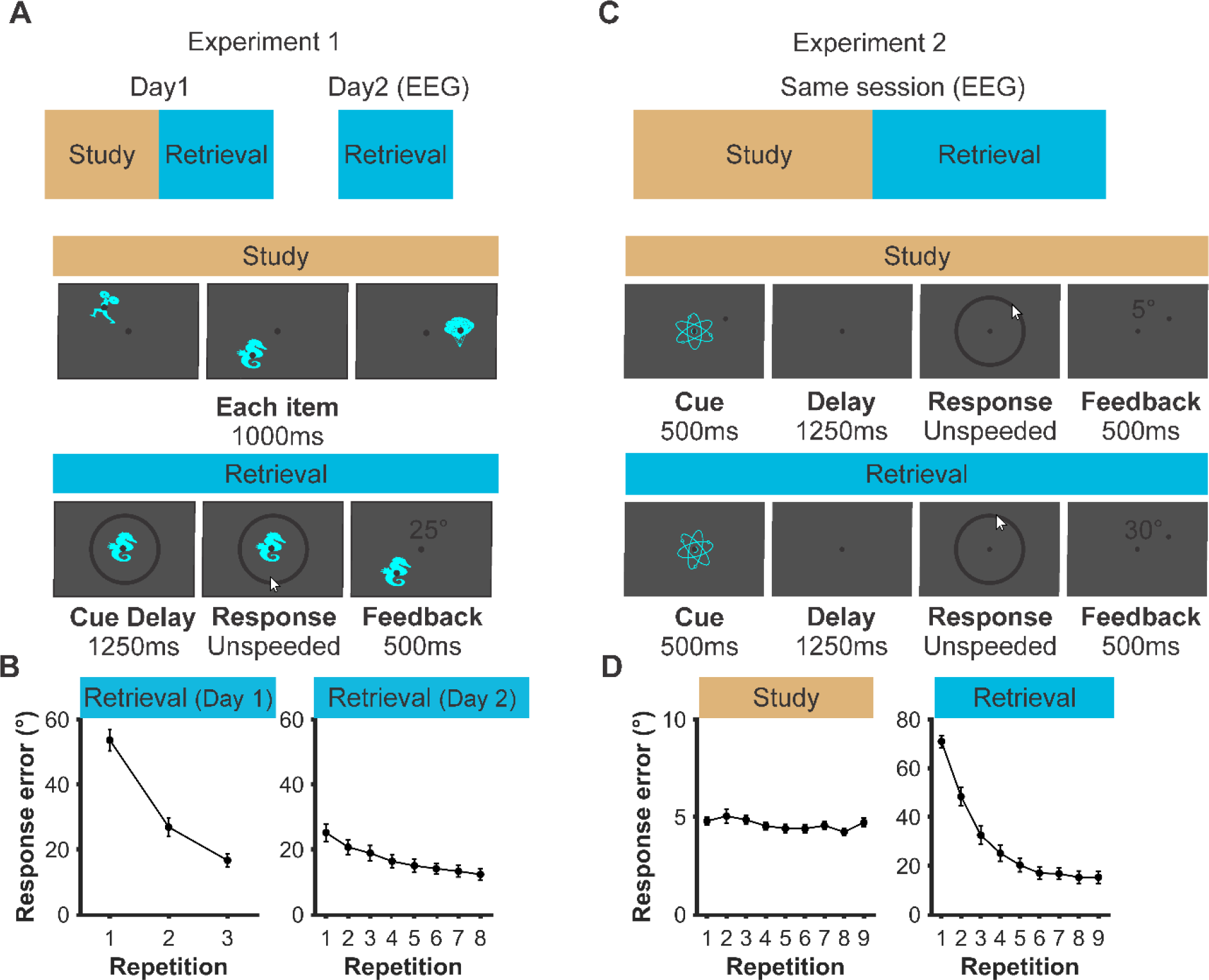
Task figure and memory performance for Experiments 1 and 2. **A.** Schematic of Experiment *1,* with example study (Experiment 1, Day 1) and retrieval (Experiment 1, Day 2) trials. Each trial was initiated with a space press. **B.** Average absolute error of retrieval responses demonstrating improvements over retrieval repetitions. **C.** Schematic of interleaved study and retrieval repetitions in Experiment 2 with exam pie study and retrieval trials. Each trial was initiated with a space press. **D.** Average absolute error of study and retrieval responses demonstrating memory accuracy over study and retrieval repetitions. Error bars represent ±1 s.e.m.

On Day 1, all 120 object-location pairings were studied over three repetitions with interleaved retrieval practice. Each of these repetitions were randomly divided into 12 “mini-blocks”, in which 10 objects were presented followed by a final test on all objects in a random order. Specifically, 10 objects were serially presented in their respective spatial locations (1000 ms per object, each object initiated by pressing spacebar). Next, each of the 10 objects were presented at fixation in a random order (1000 ms per object), and participants clicked that object’s location along a ring (unspeeded). Recall performance was assessed by calculating the response error (i.e., difference between the presented and reported location, ranging between –180° and 180°). After each response, participants were presented with the object in its correct location and the response error (500 ms). After completing these mini-blocks, participants again retrieved all 120 objects in a random order. One participant did not complete the final retrieval on the third run, and one participant accidentally aborted the experiment during the presentation of the first 10 objects before completing the rest of the session.

On Day 2, participants repeatedly retrieved the location of all 120 objects while we recorded EEG activity (Figure 1a). During each repetition (7–8 in total), the objects were presented in a random order. Each retrieval trial was initiated by a space press. After a variable interval of 1100 to 1500ms, an object was presented at fixation along with the response ring. Participants were instructed to maintain fixation and to avoid blinking or moving the mouse from trial initiation until the cursor appeared. Participants were also instructed to recall the location during the retrieval delay (1250 ms).

**Experiment 2 task procedure.** Experiment 2 was designed to examine encoding-retrieval similarity within a single session. As such, the experiment was modified to take place within one day by reducing the total number of objects (80 vs. 120). Participants were instructed to learn object-location associations as accurately as possible and that they would alternate between studying and being tested on these associations (Figure 1c).

During the study session, participants studied and then recalled each item during each trial. Each study trial was initiated by a space press. After a variable interval of 500 to 800 ms, an object was centrally presented (Figure 1c) along with a dot at the paired location (500 ms stimuli) followed by a blank delay (1250 ms). To prevent participants from using a part of the object as a reference to remember the associated location, we randomly varied the orientation of each object (−45 to 45°) for each presentation. As in Experiment 1, participants then reported the to-be-remembered location by clicking on the response ring (unspeeded). Participants were instructed to click with the left mouse button if they were confident in their response, and to click with the right mouse button if they felt that they were guessing. Both confident and guess responses were used for subsequent analyses. After each response, participants were shown the correct location of the item along with a number denoting the magnitude of the error. After studying all 80 objects, participants underwent another retrieval test for all objects in a random order (Figure 1c). The only difference between study and recall trials was the presence of the peripheral dot.

**Stimuli.** In Experiment 1, 120 clip art objects (e.g., animals, plants, objects) were selected from the Sutterer and Awh (2016) clip art library. All objects were randomly assigned to unique angular locations (0–360°, 3° steps) for each participant. On Day 1, the viewing distance was ∼80 cm (1.9° stimuli, 5° response ring, 0.3° fixation dot). On Day 2, the viewing distance was ∼100cm (1.5° stimuli, 4° response ring, 0.25° fixation dot). The background of the screen was medium gray, all objects appeared in the color cyan, the response ring was dark grey, and the fixation dot was rendered in black.

In Experiment 2, 80 of the objects from Experiment 1 were randomly paired with a unique location drawn from all 360° of possible locations. In order to assure that the entire space was used, assignment of locations was constrained such that an equal number of locations were drawn without replacement from eight bins each spanning 45 degrees of the possible space. The viewing distance was ∼100 cm (1.2° stimuli, 4° response ring, 0.25° fixation dot). The background of the screen was again medium gray, all objects appeared in the color cyan, and the response ring and the fixation dot were dark grey.

**Modeling of Response Errors.** Response error was measured as the number of degrees between the presented angular location and the reported angular location. Errors ranged from 0° (a perfect response) to ±180° (a maximally imprecise response). For each run (Figure 1b), we calculated the average absolute response error for the artifact free trials. Error distributions of this sort have been shown to be well described by a mixture of a uniform distribution for guesses and a Von Mises distribution for correct responses (Zhang and Luck 2008; Brady et al. 2013). We used MemToolbox (Suchow et al. 2013) to calculate the probability of retrieval (*P_mem_*), precision (*SD*), and the bias (*μ*) of each participants responses.

**EEG acquisition.** In Experiment 1, EEG was recorded with 20 tin electrodes mounted in an elastic cap (Electro-Cap International, Eaton, OH). We recorded from International 10/20 sites F3, FZ, F4, T3, C3, CZ, C4, T4, P3, PZ, P4, T5, T6, O1, and O2, along with five nonstandard sites (OL, OR, PO3, PO4, POz). All sites were recorded with a left-mastoid reference, and were re-referenced offline to the algebraic average of the left and right mastoids. To detect horizontal eye movements, electrodes were placed ∼1 cm from the canthi of each eye to record horizontal electrooculorgram (EOG). To detect blinks and vertical eye movements, a single electrode was placed under the center of the right eye and referenced to the left mastoid to record vertical EOG. The EEG and EOG data were amplified with an SA Instrumentation amplifier, filtered (0.01–80 Hz), and digitized (250 Hz) using LabVIEW 6.1 running on a PC.

In Experiment 2, EEG was recorded from 30 active Ag/AgCl electrodes (Brain Products actiCHamp, Munich, Germany) mounted in an elastic cap positioned according to the International 10-20 system Fp1, Fp2, F7, F3, F4, F8, Fz, FC5, FC6, FC1, FC2, C3, C4, Cz, CP5, CP6, CP1, CP2, P7, P8, P3, P4, Pz, PO7, PO8, PO3, PO4, O1, O2, Oz. A ground electrode was placed in the elastic cap at position FPz. Data were referenced online to the right mastoid and re-referenced offline to the algebraic average of the left and right mastoids. Incoming data were filtered (0.01– 80 Hz) and recorded with a 500 Hz sampling rate using BrainVision Recorder running on a PC. To detect eye movements and blinks, we used eye tracking to monitor gaze position and electrooculogram (EOG) activity recorded with five electrodes (∼1cm from the outer canthi of each eye, above/below the right eye, and a ground electrode placed on the left cheek).

**Artifact Rejection.** Data from both experiments were visually inspected for EOG and EEG artifacts. Trials containing blinks, eye movements, blocking, and muscle artifacts were excluded from analysis. One electrode for one participant in Experiment 2 was also rejected during recording because it had malfunctioned. We also monitored gaze position during Experiment 2 using a desk-mounted infrared eye tracking system (EyeLink 1000 Plus, SR Research, Ontario, Canada). Gaze position data for Experiment 2 were also visually inspected for ocular artifacts. For the analysis of gaze position, we further excluded trials in which the eye tracker was unable to detect the pupil, operationalized as any trial in which the horizontal gaze position was more than 15° from fixation or the vertical gaze position was more than 8.5° from fixation. We collected useable gaze position data (500 Hz sampling rate) for 18 of 24 participants.

Removal of trials with ocular artifacts was effective: maximum variation in grand-averaged HEOG waveforms by remembered location bin was < 2.5 μV for Experiment 1 and < 2 μV for both the encoding and retrieval in Experiment 2. Thus, eye movements in both experiments corresponded to variations in eye position of < 0.2° of visual angle (Lins et al. 1993), roughly the size of the fixation dot. Analysis of the subset of participants (18) for whom we were able to obtain reliable gaze position data in Experiment 2 corroborates the HEOG data obtained from all participants. Variation in grand-average horizontal gaze position as a function of remembered location was < 0.11° for encoding and < 0.08° of visual angle for retrieval. Variation in grand-average vertical gaze position data by remembered location was < 0.14° for encoding and < 0.09° of visual angle for retrieval. For comparison, HEOG for these participants showed a < 2.1 μV maximum variation which also corresponds to < 0.2° of visual angle.

**Time-frequency analysis.** To calculate frequency specific activity at each electrode we first band-pass filtered the raw EEG data using EEGLAB (eegfilt, see Delorme and Makeig, 2004). Alpha band analyses were band-pass filtered between 8 to 12 Hz, which is consistent with our prior work (Foster et al. 2016). For our exploratory analysis of the full range of frequencies, we band-pass filtered the data at 1 Hz intervals (4–50 Hz, down-sampled to 20 Hz, filter order:

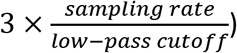

We then applied a Hilbert transform (MATLAB Signal Processing Toolbox) and squared the complex magnitude of the complex analytic signal for each trial to calculate instantaneous power before averaging across trials.

**Inverted encoding model.** Following our prior work (Foster et al., 2016), we reconstructed spatially selective channel-tuning functions (CTFs) from the multivariate topographic distribution of oscillatory power across electrodes. We assumed that the power at each electrode reflects the weighted sum of eight spatially selective channels (which we assume reflect the responses of neuronal populations), each tuned for a different angular location (Brouwer and Heeger 2009; Sprague and Serences 2013; for review, Sprague et al. 2015; Foster et al. 2016). We modeled the response profile of each spatial channel across angular locations as a half sinusoid raised to the seventh power:

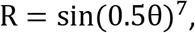

where θ is angular location (0–359°), and *R* is the response of the spatial channel in arbitrary units. This response profile was circularly shifted for each channel such that the peak response of each spatial channel was centered over one of the eight location bins. These 8 location bins each spanned 45° and were centered on 22.5°, 67.5°, 112.5° etc for Experiment 1 and on 0°, 45°, 90° etc for Experiment 2. Bin centers for each experiment were chosen prior to data collection.

An IEM routine was applied to each time point in the alpha-band analyses and to each time-frequency point in the time-frequency analyses. We partitioned our data into independent sets of training data and test data (for details see the Assigning trials to training and test sets section). This routine proceeded in two stages (train and test). In the training stage, training data *B_1_* were used to estimate weights that approximate the relative contribution of the eight spatial channels to the observed response measured at each electrode. Let *B_1_* (*m* electrodes × *n_1_* observations) be the power at each electrode for each measurement in the training set, *C_1_* (*k* channels × *n_1_* measurements) be the predicted response of each spatial channel (determined by the basis functions) for each measurement, and *W* (*m* electrodes × *k* channels) be a weight matrix that characterizes a linear mapping from “channel space” to “electrode space”. The relationship between *B_1_*, *C_1_*, and *W* can be described by a general linear model of the form:

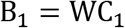

The weight matrix was obtained via least-squares estimation as follows:

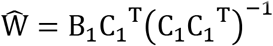

In the test stage we inverted the model to transform the observed test data *B_2_* (*m* electrodes × *n_2_* observations) into estimated channel responses, *C_2_* (*k* channels × *n_2_* measurements), using the estimated weight matrix, 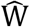, that we obtained in the training phase:

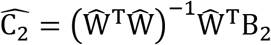

Each estimated channel response function was then circularly shifted to a common center (i.e., 0° on the “Channel Offset” axis of Figure 2a) by aligning the estimated channel responses to the channel tuned for the cued/target location to yield the CTF averaged across the eight remembered locations.

**Figure 2.**
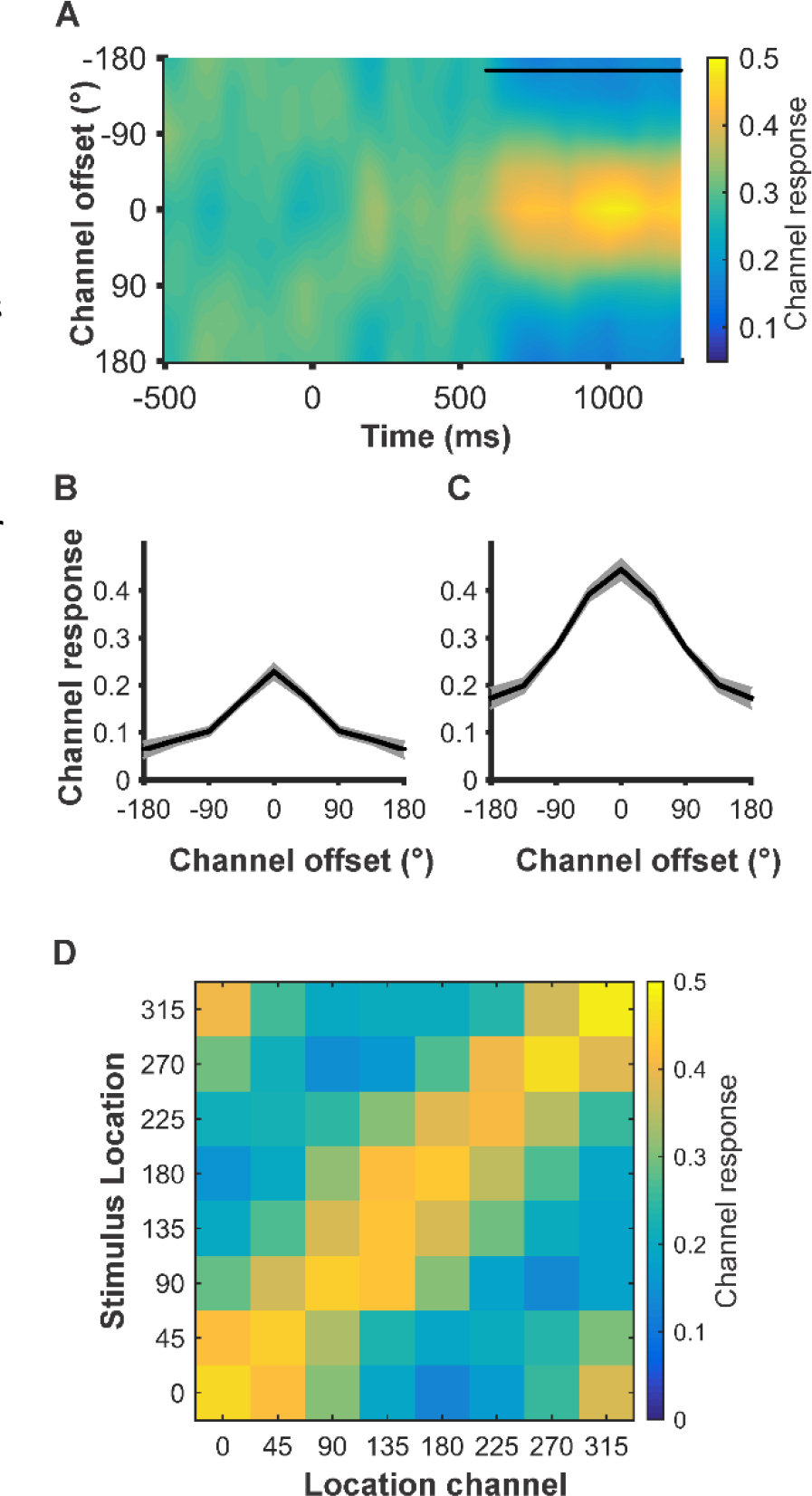
Alpha band (8–12 Hz] channel tuning functions (CTF)from Experiment 1. **A.** Alpha CTF across time. An I EM was used to reconstruct spatially selective CTFs from the topographic distribution of alpha-band power. CTF selectivity was reliable from 588 to 1250ms (quantified as CTF slope; p < .05, indicated by the black marker). B. Alpha CTF derived with a set of 8 delta functions and averaged across significant time points (588–1250 ms). Delta function CTFs are graded confirming that the signal carried by the topography of alpha band power is Intrinsically graded. Thus, the use of a graded basis set is appropriate. Shaded area represents ±1 bootstrapped s.e.m. **C.** Alpha CTF derived with a graded basis set and averaged across significant time points (588–1250 ms). Shaded area represents ±1 bootstrapped s.e.m. **D.** Channel responses for each of the eight stimulus location bins averaged across significant time points (588–1250ms), The channel response peaks at the channels preferred location, indicating that alpha activity is selective for the specific remembered location.

Finally, because the exact contributions of each spatial channel to each electrode (i.e., the channel weights, W) varies across participants, we applied the IEM routine separately for each participant, and statistical analyses were performed on the reconstructed CTFs. This approach allowed us to disregard differences in the how location-selective activity is mapped to scalp-distributed patterns of power across participants, and instead focus on the profile of activity in the common stimulus or “information” space (Sprague et al. 2015; Foster et al. 2016; Foster, Sutterer, et al. 2017).

**Assignment of trials to training and test sets.** Artifact free trials were partitioned equally into three independent sets to be used as training and test data for the IEM procedure (see Inverted Encoding Model). We down-sampled the data so that each set contained an equal number of trials, and that each location bin within a set also contained the same number of trials. For each of these sets we averaged power across trials for each location bin. We used a cross validation routine such that two sets of estimated power served as the training data and the remaining set served as the test data. We applied the IEM routine using each of the three matrices as test data, and the remaining two matrices as training data. The resulting CTFs were averaged across each test set.

For the analysis in which we ruled out the possibility that the IEM was detecting object-specific information, we assigned all trials with the same object to the same partition. After completing this additional step, we equated trials across sets and bins in the same manner described above.

For analyses in which we examined how within participant changes in selectivity related to behavior, we first down-sampled to equate the number of trials assigned from each location across conditions. After completing this additional step, we equated trials across sets and bins in the same manner described above. Finally, we employed the same training procedure described above (2/3 of the total data), but split the final test set into our comparisons of interest. Thus, we used the same training data for both conditions and only the test data varied for each comparison.

For analyses that assessed relationships between CTF selectivity and behavior across participants we down-sampled the number of trials assigned to each location bin for each of the three sets to be equal to the smallest number of trials assigned to each bin in each set for any participant. This down sampling approach precluded individual differences in CTF selectivity driven by the number of the trials included in the analysis for each participant.

In Experiment 2, we sought to compare encoding and retrieval related activity. We closely followed the procedure that examined retrieval-related activity alone, by training on 2/3 of the encoding data and testing on 1/3 of the retrieval data. By maintaining these same ratios of training to test data, we could more directly compare the results from encoding and retrieval.

**Resampling random assignment.** To avoid spurious results due to the random assignment of trials, we repeated each analysis multiple times with a different random assignment of trials. When comparing between conditions, we conducted 500 iterations per time point. When comparing against a permuted null distribution (which is a time consuming procedure), we conducted 10 iterations per time point, given the computational time needed for each analysis. In order to decrease computation time further for the 4–50 Hz time-frequency analysis, we down sampled the data matrix of power values to 50 Hz (i.e., one sample every 20 ms). We down sampled after calculating power so that down sampling did not affect our calculation of power. The data matrix was not down-sampled for analyses restricted to the alpha band.

**Calculating CTF Selectivity.** To quantify selectivity at each time point we calculated the slope of the CTF via linear regression. We collapsed across channels of equidistance (e.g., ±2 bins). As such, higher slope values indicate greater CTF selectivity while lower values indicate less CTF selectivity.

To test whether CTF selectivity was reliably above chance, we tested whether CTF slope was greater than zero using a one-sample *t* test. Because mean CTF slope may not be normally distributed under the null hypothesis, we employed a Monte Carlo randomization procedure to empirically approximate the null distribution of the *t* statistic. To generate our null distribution, we randomly shuffled the remembered location labels in each training/test set so that the labels were random with respect to the observed responses at each electrode. We then repeated 1000 iterations of this randomization procedure to obtain a null distribution of *t* statistics at each time point.

Finally, to test whether CTF selectivity was reliably above chance we employed a nonparametric cluster approach that corrects for multiple comparisons by taking into account auto-correlation in time and frequency (Maris and Oostenveld 2007; Cohen 2014). Specifically, we applied a t-value threshold corresponding to *p* < .05 (Experiment 1: *t* = 1.706; Experiment 2: *t* = 1.714) to identify clusters of pixels (time and frequency analysis) or adjacent time points (alpha only analysis). At the same time, we applied the same threshold to each permutation and calculated the largest summed-t statistic for any cluster in the permutation, resulting in a distribution of maximal summed t-statistics for our permuted null distribution. Finally, the sizes of the significant clusters of the non-permuted data were thresholded such that only clusters larger than the 95^th^ percentile of the permuted distribution were considered reliable (Type 1 error less than .05). Therefore, our cluster test was a one-tailed test, corrected for multiple comparisons.

**Resampling test.** When comparing between conditions (i.e., baseline alpha power, CTF slope), we used a non-parametric resampling procedure across participants (Efron and Tibshirani 1993). We resampled each participant with replacement 100,000 times. Then, we calculated the number of these resampling iterations in which the differences were ≤ 0 (one-tailed). In cases where no iterations were ≤ 0, we report the p values as *p* < .001. We deemed results to be reliably above chance if *p* < .05.

## Results

### Experiment 1

Experiment 1 was designed to test whether continuous tests of memory accuracy, in conjunction with an IEM applied to EEG data, could be used to track memory retrieval. The design includes two important properties. First, we used a continuous test of memory accuracy by having participants report remembered locations along a ring. This provides a sensitive test of memory accuracy as the deviation from the correct location. Second, we recorded EEG activity during memory retrieval for the purposes of building and evaluating an encoding model. This inverted encoding model (IEM) can track memory retrieval as a graded function of spatial location.

**Behavioral performance.** On Day 1, participants studied 120 object-location associations (Figure 1a). On Day 2, participants returned for a retrieval session in which we recorded EEG. Participants received feedback based on their response error (−180° to 180°). During both days, their performance improved (Figure 1b). During Day 1, average response error improved significantly from the first (*M* = 53.6°, *SD* = 17.7°) to the final test (*M* = 17.6°, *SD* = 11.1°) of the session (*t*(26) = 13.2, *p* <.001, one-tailed). As a result on continued feedback, memory also improved from the first (*M* = 25.2°, *SD* = 14.1°) to the final test (*M* = 12.7°, *SD* = 9.3°) during the second session (*t*(26) = 7.3, *p* <.001, one-tailed).

**Alpha-band (8–12 Hz) topography tracks spatial representations retrieved from LTM.** In Experiment 1, we tested whether oscillatory EEG activity tracks the time-resolved retrieval of precise spatial memories. Because we have previously found that alpha-band activity tracks spatial locations held in working memory (Foster et al. 2016), we were *a priori* interested in whether alpha-band power would also track locations retrieved from long-term memory. Thus, we used an IEM to test whether the multivariate topography of alpha-band power tracked locations retrieved from long-term memory. If the pattern of alpha-band power contains spatially selective information about the remembered location, we would expect to see a graded channel tuning function (CTF) with a peak response in the channel tuned for the remembered location (a channel offset of 0°in Figure 2) following the retrieval cue. This graded pattern can be quantified as slope across the position channels as distance from the retrieved location increases. A slope of zero reflects no spatial selectivity in the CTF, while a positive slope reflects spatial selectivity for the location associated with the cue. To test this hypothesis, we conducted a permutation test (see Materials and Methods) to determine at which time points we observed a CTF slope that was reliably above zero. We detected reliable selectivity for spatial information (i.e., slopes > 0) that was sustained during the retrieval interval (588 – 1250ms; Figure 2a).

One possibility is that this graded tuning is an artifact of our selection of a graded basis set (Saproo and Serences 2014; Ester et al. 2015; Foster et al. 2016). To investigate this possibility, we reran this analysis using a delta function basis set that predicts a peak response in the preferred channel and no response in adjacent channels. If the topography of alpha power represents spatial locations in a graded manner, we would still expect a graded pattern of responses. Instead, if the observed results were driven by our selection of a basis set, we would expect a peak response in the correct bin and a little to no response in all other bins. Using a delta function basis set, we observed a graded pattern of responses across remembered locations (Figure 2b) that is similar to the pattern of activity we see when we apply the standard basis set (Figure 2c). This suggests that our results are not an artifact of our selection of a basis set, but reflect a real graded tuning profile during the retrieval of spatial memories.

Although the aggregate results revealed that channel activity peaked at the remembered location and dropped in a graded fashion as the distance form that location increased, this analysis did not establish that this orderly pattern was present at each location. Indeed, a courser hemi-field or quadrant-based signal could produce such a pattern. If alpha-band activity precisely tracks retrieved spatial locations, we should observe a graded pattern for each remembered location. We examined the average channel response during time points where we previously observed reliable spatial selectivity (588–1250ms) for eight location bins separately.

The channel response for each location revealed graded information throughout the same window, and the channel response for all locations were reliable (All slopes > 0.05, all *p’s* < .002). Therefore, alpha-band CTFs track memory retrieval of a precise spatial location.

**Identifying frequencies that track the retrieval of spatial location.** A motivating question for the present work was whether spatially selective information was specific to alpha-band activity. On the one hand, prior work has found that alpha-band activity selectively tracks spatial locations that are covertly attended (Foster, Sutterer, et al. 2017) or held in working memory (Foster et al. 2016). On the other hand, theta-band (4–7 Hz) and beta-band (16–25 Hz) activity are known to play an important role in long-term memory (Nyhus and Curran 2010; Morton et al. 2013). Therefore, we performed the same IEM analysis at each frequency and time point from 4–50 Hz to test whether other frequency bands also carried spatially selective information about the remembered location. We conducted a permutation test at each frequency and time point and used a cluster correction to identify frequencies with CTFs that were reliably above zero (Figure 3a). Although we observed brief periods of spatial selectivity in the beta range (16–25 Hz) the most robust and sustained selectivity was in the alpha band (410ms – 1250ms).

**Fig 3.**
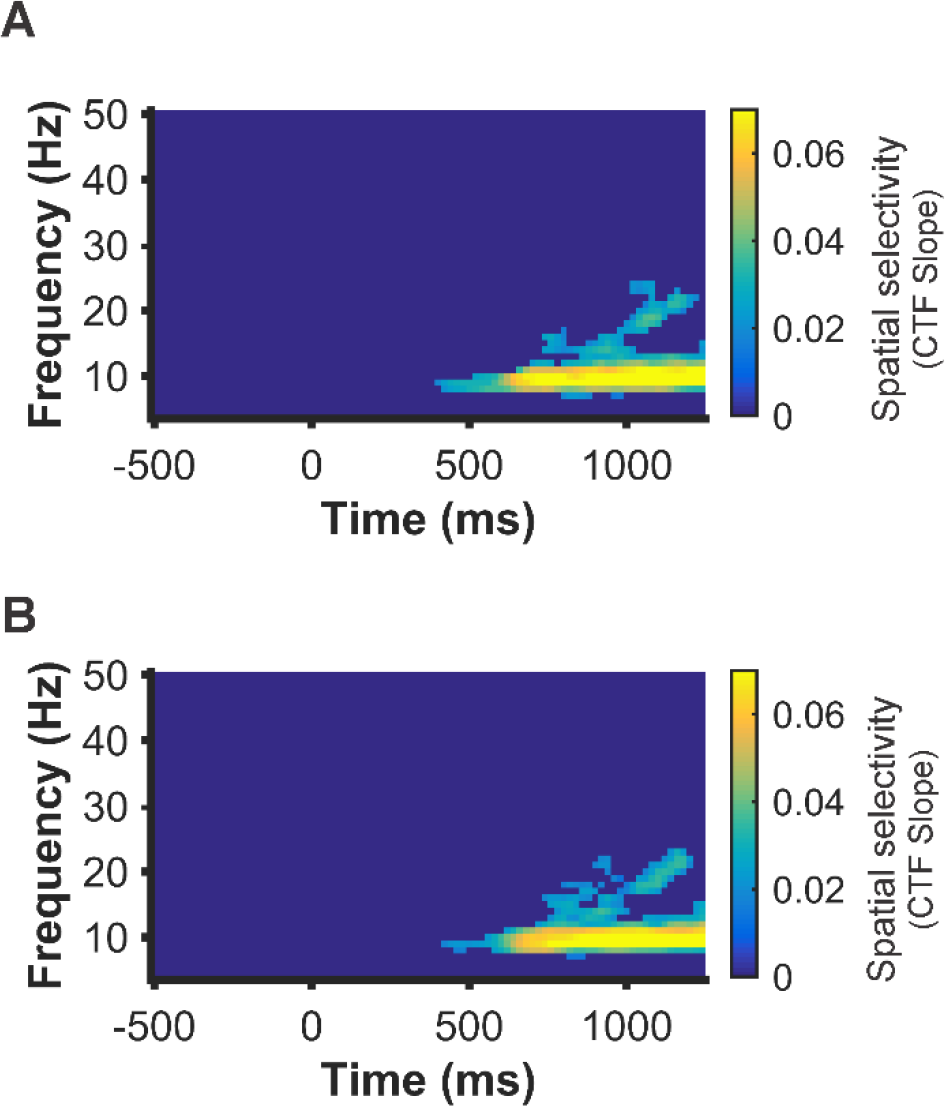
Identifying frequencies that track retrieved locations for Experiment 1. **A.** An IEM was used to reconstruct spatially selective CTFs from the topographic distribution of total power across a range of frequencies (4–50Hz). Alpha and to a lesser extent beta power tracked retrieval of spatial information. **B.** Training and testing across shapes. Alpha power continued to track the retrieval of spatial information when the IEM was trained and tested on separate shape cues, indicating that CTFs reflect remembered locations not the retrieval cue. Points at which CTF slope values were not reliably above zero as determined by a cluster corrected permutation test (p <.05) were set to dark blue.

**Spatially selective alpha-band activity generalizes across visual objects associated with the same spatial location.** For each participant, each object was associated with a unique location such that object and position were confounded within this analysis. Thus, it is possible that the selectivity we observed across some or all frequencies, reflects patterns of activity elicited by the cue rather than activity related to the retrieval of a spatial position. To investigate this, we re-ran the analysis while ensuring that distinct items were included in the training and test sets (see Materials and Methods). Despite this constraint, we observed similar results (Figure 3b), confirming that the sustained spatial selectivity we observed reflected the position associated with each cue rather than the cue itself.

**Spatially selective alpha-band activity tracks the accuracy of recall from long-term memory**. A consequence of providing feedback during Day 2 is that memory performance improved throughout the session. To examine whether alpha-band CTFs tracked these behavioral improvements, we split the test data into the first half and second half of trials. Behaviorally, we observed that memory performance improved from the first half (*M* =20.3°, *SD* = 11.9°) to the second half (*M* = 13.9°, *SD* = 9.4°) of the session (*t*(26) = 7.2, *p* <.001, one-tailed, Figure 4a). Furthermore, a mixture model was used to assess whether these decreases in average response error were driven by changes in the probability of retrieval and/or mnemonic precision (see Materials and Methods). During Day 2, the probability of retrieval (*P_mem_*) increased over time (first half: *M* = 86.3%, *SD* = 13.6%; second half: *M* = 93.6%, *SD* = 9.5%; *t*(26) = −6.3, *p* <.001, one-tailed) and mnemonic precision improved over time (first half: *M* = 13.6°, *SD* = 3.7°; second half: *M* = 12.3°, *SD* = 4.1°, *t*(26) = 5.52 *p* <.001, one-tailed). If alpha-band CTFs are sensitive to the accuracy of memory retrieval, we would expect greater spatial selectivity in the second vs. first half of the session. To test this prediction, we used a resampling test (see Materials and Methods) in which we isolated the time points where aggregate data revealed significant alpha CTFs (Fig 2a; 588ms after cue onset until the response), and then compared average CTF slope across the first and second halves of the study. Indeed, spatial selectivity was significantly higher for the second half (CTF slope, *M* = 0.085, *SD* = 0.061) relative to the first half (*M* = 0.06, *SD* = 0.055) of the experiment (Figure 4b) (p <.001, one-tailed resampling test).

**Figure 4.**
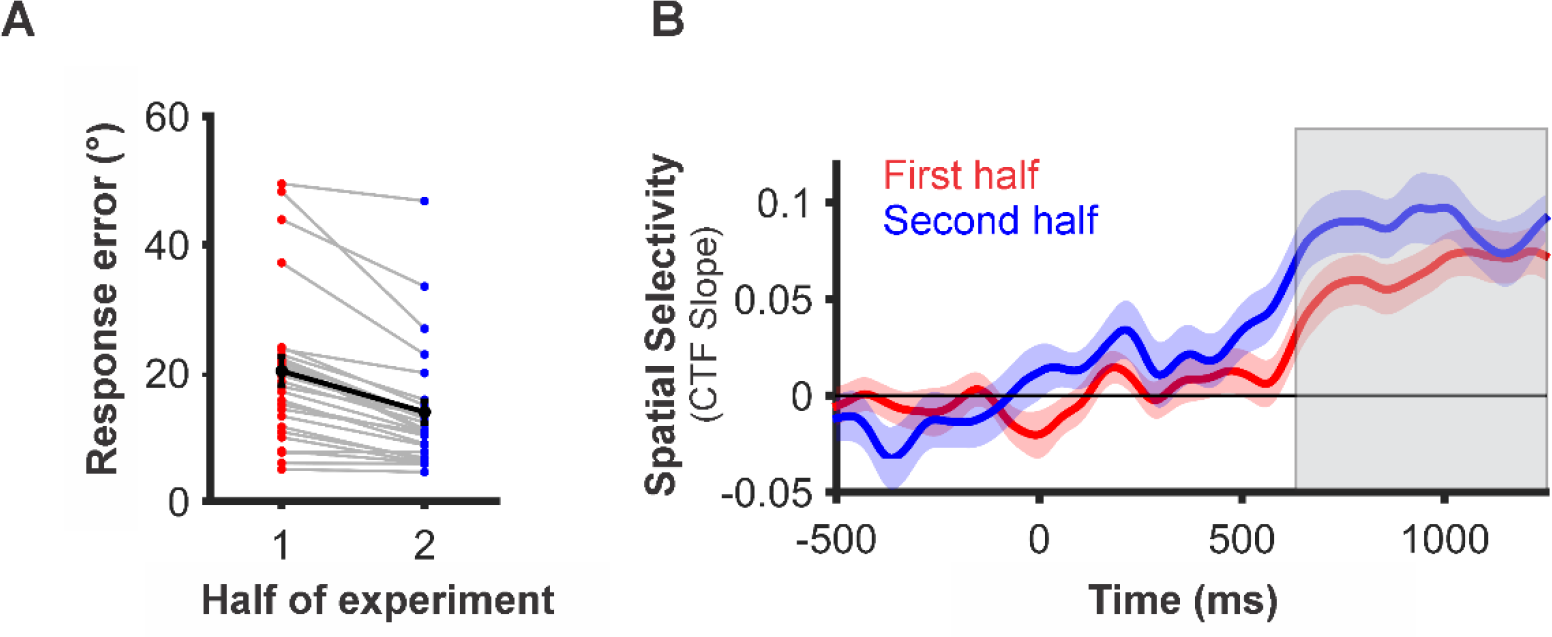
Assessing the relationship between alpha selectivity and memory performance for Experiment 1. **A.** Average response error improved from the first to the second half of the experiment (p<.001). Error bars represent ±1 s.e.m. **B.** Time resolved CTF slopes for trials from the first and second half of the experiment. CTF slope reflects learning across the session and reveals significantly higher spatial selectivity for the second half of the experiment relative to the first half (p<.001). Reliable differences were assessed by averaging across time points where we observed reliable CTFs for all trials (grey box) and comparing CTF slope between the first and second half of the experiment. Shaded error bars represent ±1 bootstrapped s.e.m.

This reveals that alpha-band activity tracks the improvement in memory performance across learning episodes. Finally, CTF selectivity across the same window did not predict between-subject variations in the accuracy of recall (*ρ*(26) = − .11, *p* =.6). This null result could have numerous explanations but here we offer one hypothesis. While we instructed participants to immediately recall the location that corresponded to the object cue, it may be that some participants waited longer than others to call the correct location to mind while other participants relied on a more prospective strategy in which they immediately recalled the target location. This kind of strategic difference could yield large differences in mean CTF slope that may have been unrelated to whether the critical item could be retrieved. Indeed, the response time analysis in the next section lends further plausibility to this hypothesis.

**Spatially selective alpha-band activity tracks within- and between-subject variations in response latency.** The latency of memory retrieval varied across trials and participants to a large extent (see Fig 5a). To examine whether alpha-band CTFs tracked these behavioral differences in response time (RT), we split the test data into two halves based on the median RT (average fast RT: *M* = 854ms, *SD* = 240ms; average slow RT: *M* = 1961ms, *SD* = 1055ms). If alpha-band CTFs track the latency of memory retrieval, we would expect greater location selectivity on trials in which participants responded more quickly. Indeed, location selectivity was significantly greater when participants responded more quickly (*M* = 0.085, *SD* = 0.054) than when they responded slowly (*M* = 0.052, *SD* = 0.060; *p* < .001, one-tailed resampling test; Figure 5b). This pattern supports the hypothesis that participants responded more quickly when they had already retrieved the spatial information prior to the onset of the response cue, yielding a higher level of CTF selectivity during trials with faster responses.

**Figure 5.**
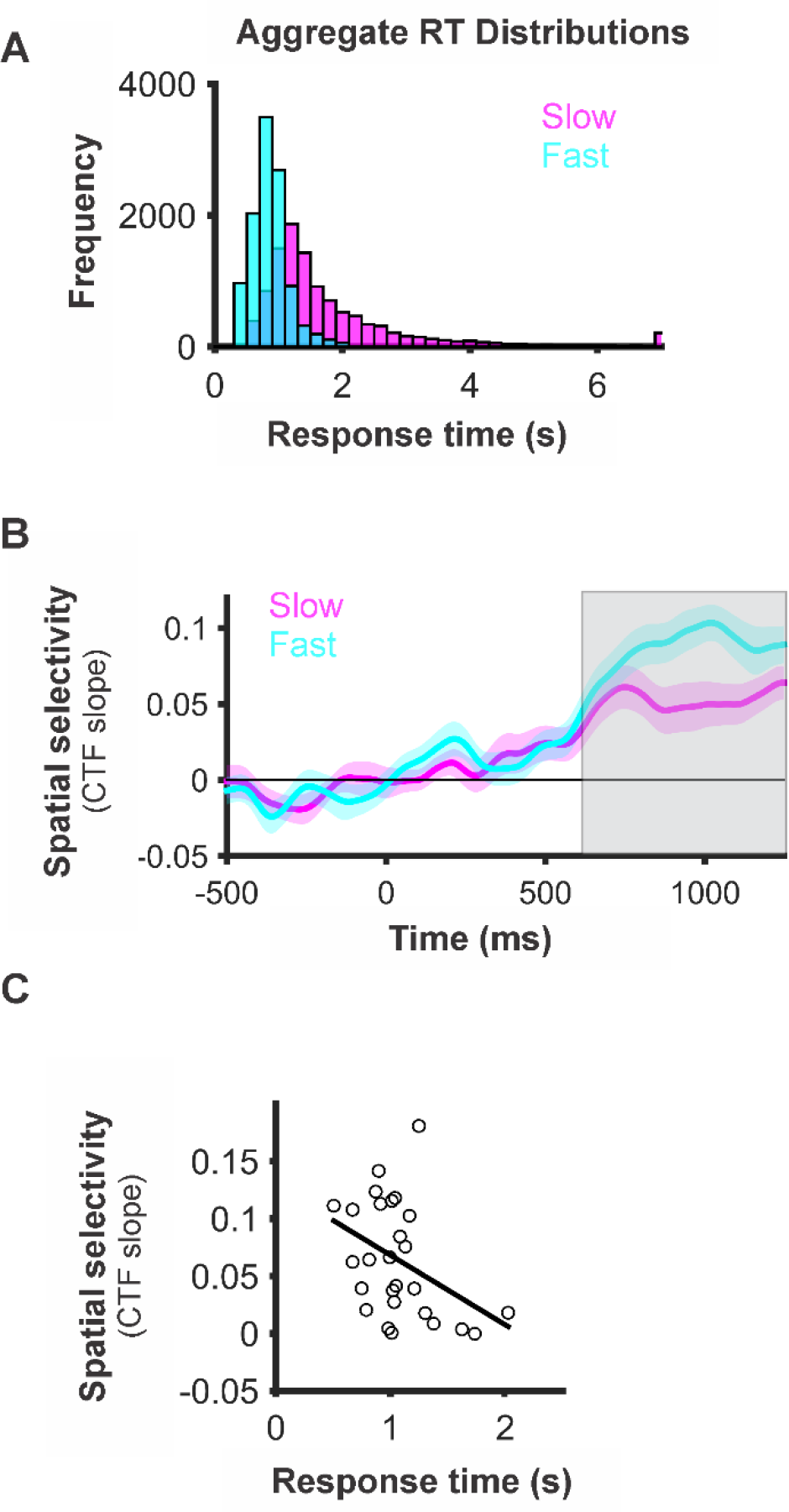
Assessing the relationship between alpha selectivity and response times for Experiment 1. **A.** Aggregate distribution of all participants’ fast and slow response times. Response times > 7s a re represented in the last bin of the histogram. **B.** Time resolved CTF slope for trials with the fastest and slowest response times. CTF slope reflects response latency and reveals that spatial selectivity was higher for trials when participants responded quickly (p<.001). Reliable differences were assessed by averaging CTF slope across time points where we observed reliable CTFs for all trials (grey box) and comparing splits with a resampling test. Shaded error bars represent ±1 s.e.m. **C.** Alpha selectivity is modestly correlated with response times across subjects although the relationship is not significant (ρ(26)= −.36 p = .07).

Across participants, we observed substantial variation in median RTs (range = 504 – 2025 ms). To examine whether alpha-band CTFs tracked these individual differences in behavior, we tested whether there was a correlation between median RT and the selectivity of alpha-band CTFs (measured as slope). We predicted that participants who responded more quickly (i.e., faster RTs) would also have greater spatial selectivity (i.e., higher CTF slope). We observed a trending negative relationship between RT and CTF slope, as predicted (*r* = −.36; *p* = .07; Figure 5c). In addition to reflecting differences in the immediate accessibility of spatial memories, this relationship could also be driven by individual differences in the extent to which participants engaged in prospective retrieval during the delay interval. This is our working hypothesis, given that the differences in response latency seemed too large to reflect differences in the immediate accessibility of the spatial memories alone.

### Experiment 2

In Experiment 2, we replicated and extended Experiment 1 in two important ways. First, to further examine the relationship between alpha-band selectivity and memory performance, we recorded EEG data throughout the learning process, including the first retrieval attempts. Second, we recorded EEG during both encoding and retrieval, which allowed us to test the extent that retrieval-related oscillatory activity resembled encoding-related oscillatory activity.

**Behavioral performance.** During a single session, participants learned 80 object-location associations (Figure 1c) with interleaved study and retrieval. During study trials, participants actively maintained the associated spatial location over a 1250 ms delay interval. During retrieval trials, participants had to retrieve the associated spatial location from long-term memory. During study trials, memory performance was very accurate and improved modestly but reliably from the first half (*M* = 4.7°, *SD* = 1°) to the second half (*M* = 4.4°, *SD* = .9°) of the session (*t*(23) = 2.42, *p* = .012, one-tailed; Figure 1d). Mixture modelling revealed that this change was due to an improvement in mnemonic precision (first half: *M* = 5.8°, *SD* = 1.2°; second half: *M* = 5.4°, *SD* = 1.1°; *t*(23) = 2.28, *p* = .016, one-tailed) while no change was observed for probability of retrieval (first half: *M* = 99.9%, *SD* = .29%; second half: *M* = 99.9%, *SD* = .16%; *t*(23) = −.81, *p* = .21, one-tailed), which was at ceiling. For the LTM retrieval trials, we observed a substantial improvement in memory performance across the session as learning progressed. Memory error decreased from the first half (*M* = 40.8°, *SD* = 14.0°) to the second half (*M* = 16.2°, *SD* =11.9°; *t*(23) = 17.0, *p* < .001, one-tailed; Figure 6a). We replicated our finding in Experiment 1 that the reduction in memory error was driven by both an increase in the probability of retrieval (first half: *M* = 61.1%, *SD* = 16.0%; second half: *M* = 89.9%, *SD* = 13.7%; *t*(23) = −15.4, *p* < .001, one-tailed) and an improvement in mnemonic precision (first half: *M* =13.8°, *SD* = 4.35°; second half: *M* = 11.3°, *SD* =2.76°; *p* <.001, one-tailed). Thus, long-term memory improved throughout the session as participants learned the object-location associations.

**Figure 6.**
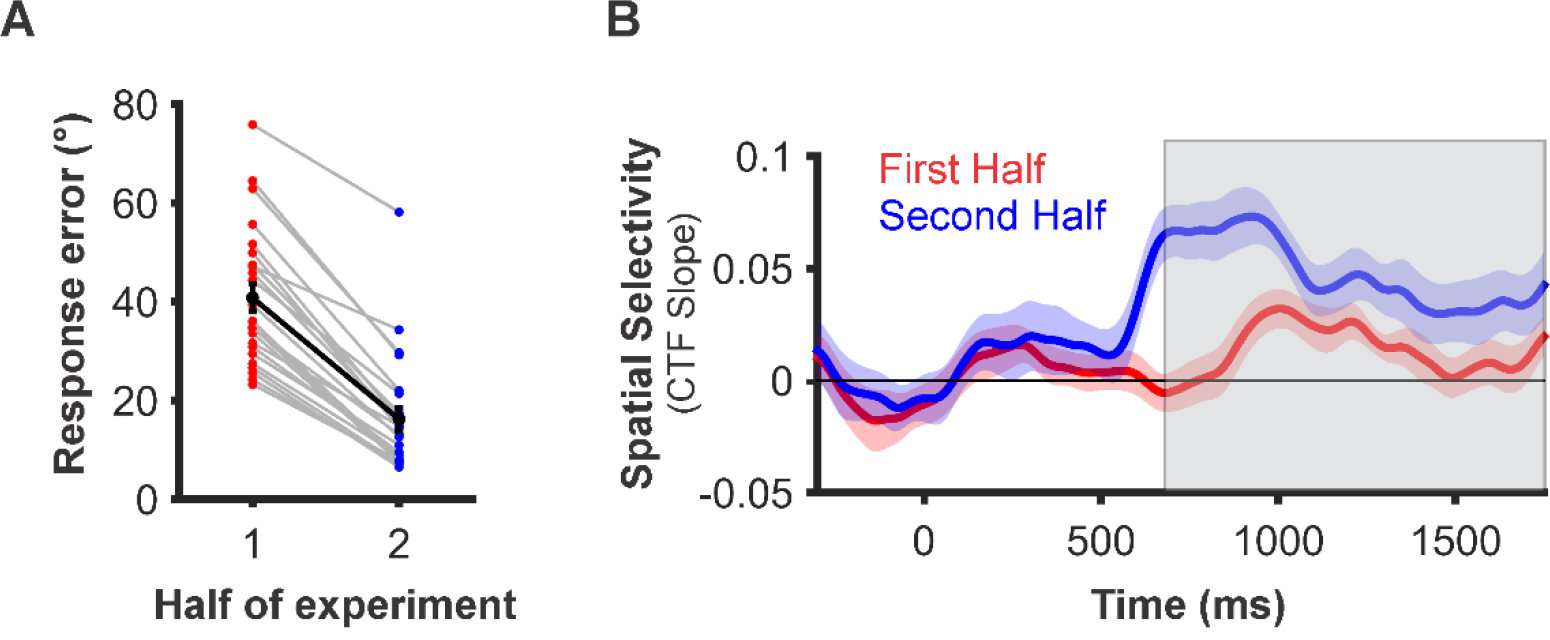
Assessing the relationship between alpha selectivity and memory performance for Experiment 2. **A.** Average response error improved from the first to the second half of the experiment (p < .001). Error bars represent ±1 s.e.m. **B.** Time resolved CTF slopes for trials from the first and second half of the experiment. CTF slope reflects learning across the session and reveals significantly higher spatial selectivity for the second half of the experiment relative to the first half (p c.001). Reliable differences were assessed by averaging across time points where we observed reliable CTFs for all trials (grey box) and comparing CTF slope between the first and second half of the experiment. Shaded error bars represent ±1 bootstrapped s.e.m.

One goal for Experiment 2 was to create a larger range of performance throughout the session in which EEG data was recorded. In line with this goal, we observed a much larger range in mean response error in Experiment 2 (71.0° Run 1 – 15.2° Run 9; Figure 1d) than in Day 2 of Experiment 1 (25.2° Run 1 – 12.3° Run 8), giving us the opportunity to apply the IEMs approach across the full trajectory of learning.

**Spatially selective alpha-band activity tracks the accuracy of recall from long-term memory**. In Experiment 1, alpha-band CTFs tracked retrieval of spatial locations from long-term memory. Furthermore, spatial selectivity of alpha-band CTFs increased as memory accuracy improved (Figure 4). Experiment 2 was designed to replicate and extend those results over a larger range of behavior. We predicted that alpha-band CTFs would demonstrate higher selectivity when memories were more accurate. In line with this prediction, the average selectivity (i.e., CTF slopes) was larger in the second half of the session (*M* = 0.048, *SD* = 0.032) than in the first half (*M* = 0.012, *SD* = 0.022; *p* < .001; one-tailed; Figure 6b). Note, for this and all subsequent average CTF analyses, we averaged from 588 ms (the starting time point used in Experiment 1) until the onset of the response cue. Control analyses revealed that increases in alpha-selectivity across the session could not be attributed to increases in alpha power across the recoding session (Figure S1). As in Experiment 1, CTF slope did not track memory performance between participants (*ρ*(23) = −.14, *p* = .52). Thus, the spatial selectivity of alpha activity tracked broad improvements in recall accuracy across the session.

**Spatially selective alpha-band activity tracks response latency.** As in Experiment 1, we found that the selectivity of alpha-band CTFs tracked within- and between-subject variations in RT (Figure 7). A median split on RT revealed greater spatial selectivity for trials with fast RTs (CTF slope: *M* = 0.035, *SD* = 0.03) than trials with slow RTs (*M* = 0.021, *SD* = 0.018), *p* = .005, one-tailed; Figure 7b). We also replicated our finding that participants with faster RTs showed greater spatial selectivity of alpha-band CTFs (*r* = −.49; *p* = .02; Figure 7c). This link between CTF slope and RTs may reflect strategic differences between participants who prospectively recalled the associated location quickly following cue onset and those that waited until closer to the response window to bring that information to mind.

**Figure 7.**
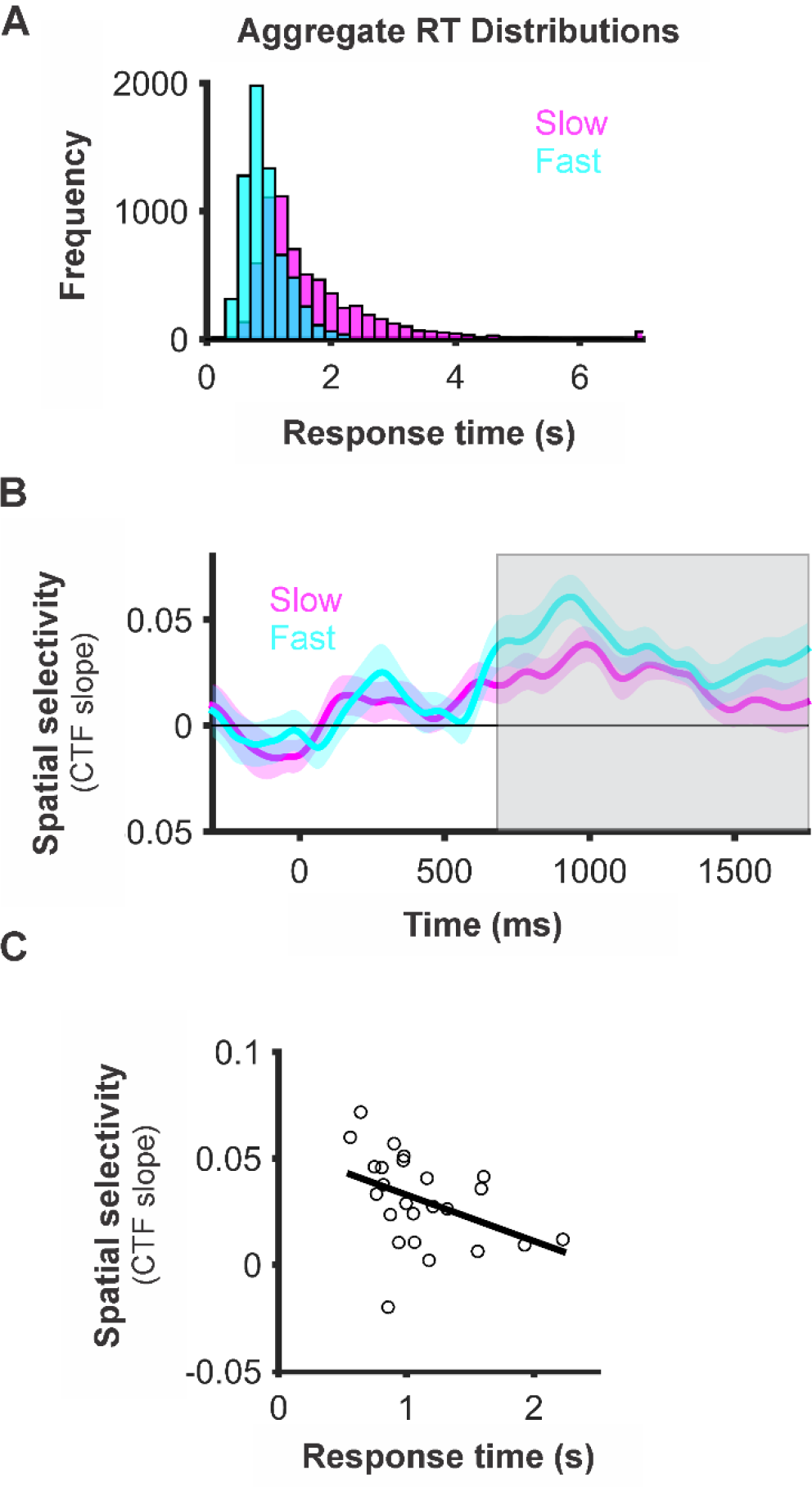
Assessing the relationship between alpha selectivity and response times for Experiment 2. **A.** Aggregate distribution of all participants’ fast and slow response times. Response times > 7s are represented in the last bin of the histogram. **B.** Time resolved CTF slope for trials with the fastest and slowest response times. CTF slope reflects response latency and reveals that spatial selectivity was higher for trials when participants responded quickly (p=.005). Reliable differences were assessed by averaging CTF slope across time points where we observed reliable CTFs (grey box) for all trials and comparing splits with a resampling test. Shaded error bars represent ±1 bootstrapped s.e.m. **C.** Alpha selectivity is modestly correlated with response times across subjects (p(23)= −.49 P = −02).

**Comparing frequency specificity at encoding and retrieval.** In Experiment 1, we found that oscillatory activity in the alpha band (8–12 Hz) tracked retrieved locations following a memory cue. In Experiment 2, we replicated this finding, with cluster corrected permutation tests showing that primarily oscillations between 8 and 12 Hz, and to a lesser extent oscillations between 12 and 18 Hz, tracked the retrieved location (∼500–1250ms; Figure 8a). Note, that in order to obtain the most robust measurement of spatially sensitive frequencies at retrieval, we only tested our IEM on trials from the second half of the experiment when memory performance and spatial selectivity were highest (Figure 6a). For consistency we applied the same approach to study trials (Figure 6b). Applying the IEM to study trials revealed that spatially selective information was represented across a wider range of low frequencies (Figure 8b; 4–8 Hz; 0–500ms; 25–30 Hz, 500–600ms; 8–20 Hz, ∼200–1750ms). Although we observed the most sustained spatial selectivity in the alpha band (8–12 Hz). These results replicate past work that has shown that alpha-band activity tracks locations held in working memory (Foster et al. 2016). Finally, an overlay plot of frequency bands carrying spatially specific information in both the encoding and retrieval tasks (Figure 8c), revealed considerable overlap in the 8–12 Hz band across conditions. Together these findings suggest that the range of frequencies carrying spatially specific information during encoding is similar to the range of spatially specific frequencies in a WM task as well as during LTM retrieval.

**Figure 8.**
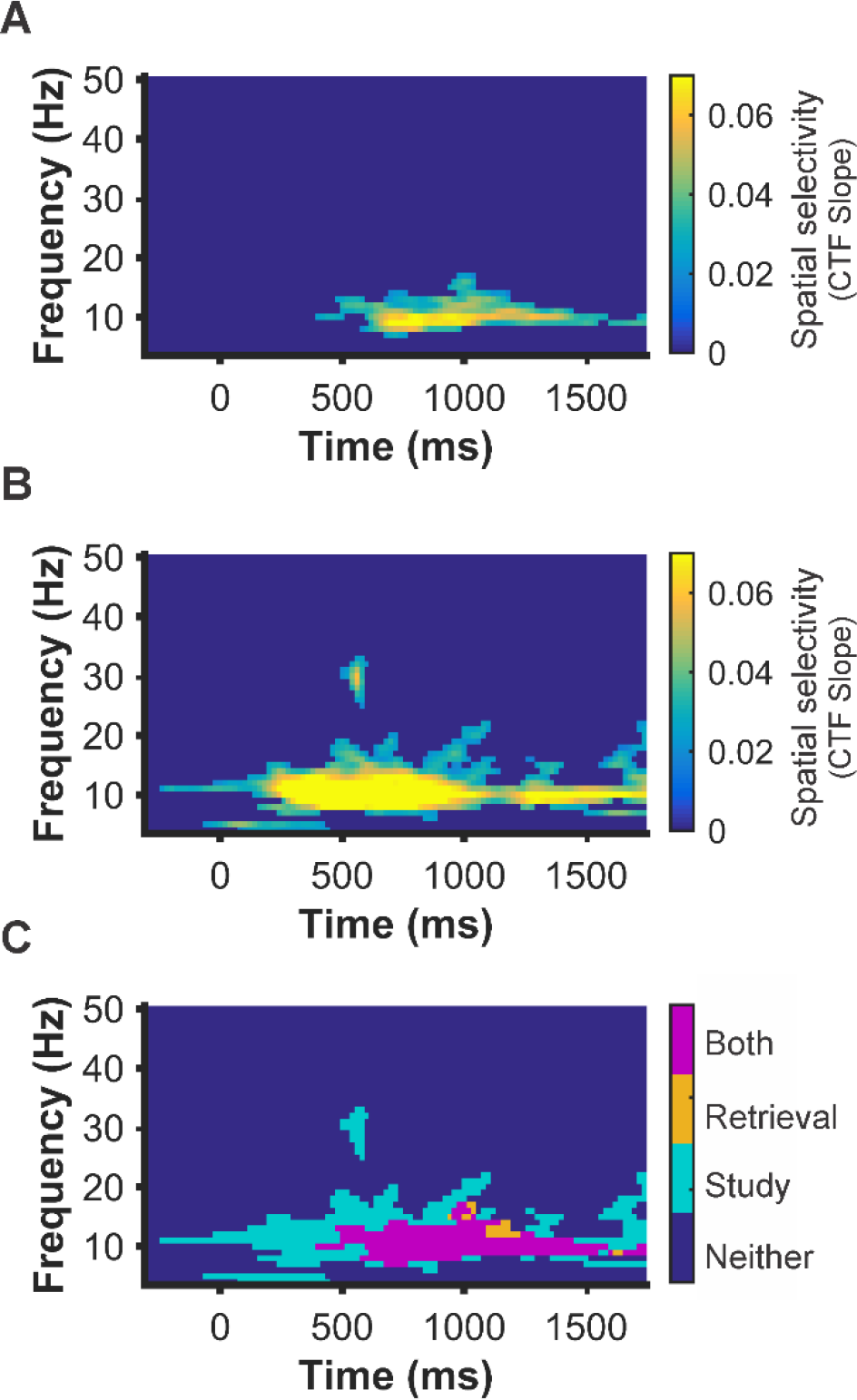
Identifying frequencies that track encoded and retrieved locations for Experiment 2. An IEM was used to reconstruct spatially selective CTFs from the topographic distribution of total power across a range of frequencies (4–50Hz) at retrieval and study. To ensure robust retrieval memory performance, only trials from the second half of the session were used In the test set for this analysis. **A.** Primarily alpha power tracked spatial information during retrieval trials. **B.** Initially, a broad range of frequencies tracked the encoded location (4-35Hz) while primarily alpha power tracked the remembered location through the entire delay. **C.** Overlay plot of significant activity during retrieval and study. Teal points reflect reliable spatial selectivity during study, orange points reflect reliable selectivity at retrieval, and magenta points reflect overlap selectivity at study and retrieval. Points at which CTF slope values were not reliably above zero as determined by a cluster corrected permutation test (p <.05) were set to dark blue.

**Patterns of alpha-band activity generalize across encoding and retrieval.** While the same frequency band carried spatially spatial information during both study and retrieval, this does not necessarily mean that the multivariate patterns of activity corresponding to each location are also similar during encoding and retrieval. To provide a comprehensive test of encoding-retrieval similarity, we trained the IEM using study trials and tested the model on retrieval trials. We observed robust spatial selectivity throughout the retrieval interval (520–1750ms; *p* < .05; Figure 9). This provides evidence that the multivariate pattern of alpha activity during retrieval is well-described as a re-instantiation of the pattern of alpha-band activity seen during encoding. The present work represents a new approach for tracking and understanding the neural mechanisms underlying retrieval of precise feature memories. Over two experiments, we employed a combination of a continuous report task, in which participants learned to associate individual objects with specific spatial locations, with the application of an inverted encoding model to ongoing EEG activity. We demonstrated that IEMs and rhythmic brain activity can be used track and reconstruct the reinstatement of spatially selective information from long-term memory.

**Figure 9.**
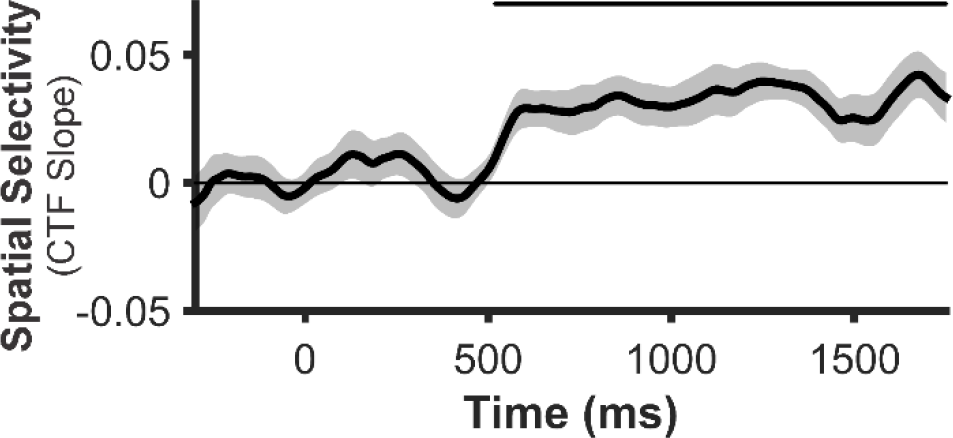
Testing whether the multivariate patterns of alpha power at study are reinstated at retrieval. Alpha power tracks the retrieval of spatial information when the IEM was trained on study data and tested on retrieval data, Indicating that the pattern of alpha band activity observed during study is reactivated at retrieval. Shaded error bars represent ±1 bootstrapped s.e.m. Markers on the top of the plot mark the periods of reliable spatial selectivity (p < .05).

We modeled long-term memory retrieval performance using a mixture modeling approach. This approach is commonplace in the field of visual working memory (Wilken and Ma 2004; Zhang and Luck 2008) but has only recently begun gaining traction in the field of long-term memory (Brady et al. 2013). This approach is able to disentangle improvements in mnemonic precision and the probability that memories are retrieved (Fan and Turk-Browne 2013; Harlow and Yonelinas 2014; Sutterer and Awh 2016). Initial studies have found that these parameters are reflected by distinct neural signals (Murray et al. 2015; Richter et al. 2016), providing further evidence that separately modeling mnemonic precision and probability of retrieval is a meaningful distinction. Our results demonstrate that both the probability of retrieving long-term memories and the precision with which those memories are retrieved continue to improve with feedback over many repetitions. We propose that this more sensitive approach of assessing memory accuracy will continue be a successful direction for the field of long-term memory.

A recent model put forth by Watrous and colleagues (2014), the spectro-contextual encoding and retrieval theory (SCERT), asserts that both the frequencies supporting cognitive operations at encoding and retrieval and the specific patterns of activity within those frequencies should overlap. In line with this prediction, we observed considerable overlap in the frequencies carrying spatial memory representations between encoding and retrieval. Furthermore, we found that the multivariate patterns of alpha-band activity reinstated during retrieval are strikingly similar to those patterns observed during the initial encoding of locations. These observations provide new evidence that encoding-retrieval oscillatory similarity extends to the representation of precise feature representations at the population level, supporting the idea that oscillatory brain activity plays a critical role in memory formation and reinstatement. However, it is worth noting that a broader range of frequencies tracked to-be-remembered locations at encoding than at retrieval, suggesting that not all spatially sensitive frequencies engaged during stimulus presentation are later reinstated.

Indeed, a novel aspect of our work was the ability to search for frequencies that code for precise spatial memories. We demonstrated robust and sustained spatial selectivity during long-term memory retrieval, primarily in the alpha band (8–12 Hz). These results are similar to what has been observed in the field of visual working memory. However this stands in contrast with other findings that suggest the role of theta and beta activity in long-term memory. In particular, past studies have found that theta activity (4–7 Hz) plays a key role in episodic memory (Nyhus and Curran 2010; Hsieh and Ranganath 2014) and in the hippocampus during spatial navigation (Watrous et al. 2011; Bohbot et al. 2017). However, it is consistent with some work that suggests a role for alpha in memory and memory guided attention (Stokes et al. 2012; Waldhauser et al. 2012, 2016). It is possible that our scalp EEG signal was relatively insensitive to theta signals prominent in the hippocampus (Hsieh and Ranganath 2014). Future work from modalities that more directly index hippocampal activity (i.e., MEG/ECOG) might provide some insight into the role of theta in precise spatial memory reinstatement.

Another promising application of the approach employed here is the ability to compare the time course with which fine-grained and courser memory representations emerge. For example, spatially selective alpha activity emerged considerably later than some prior observations of hemifield-selective activity. While hemifield selective activity has been observed within 200 ms of the onset of a retrieval cue (Waldhauser et al. 2016), our time by frequency analysis revealed no evidence of activity related to the specific retrieved location, in any frequency band, until at least 410 ms after the retrieval cue. One possible explanation for this latency difference is that hemisphere reactivation and retrieval of fine-grained spatial representations rely on different processes. For instance, Gratton et al. (1997), suggest that the hemisphere bias they observe may be more structural, resulting from the formation of a stronger trace in the hemisphere contralateral to the hemifield in which the stimulus was presented; while in the present study, the relatively slower onset of alpha CTFs implies a more effortful retrieval of precise spatial information. Another potential explanation is that context reinstatement during an object recognition task could occur more rapidly than object-cued retrieval of spatially selective information. Further work is needed to explore this difference in latency between hemifield effects and the reactivation of the fine-grained alpha topography that tracks specific locations.

Prominent models have argued that spatial-temporal context is the backbone of episodic memory (O’Keefe and Nadel 1978; Ekstrom and Ranganath 2017) serving as an index for the retrieval of specific past experiences. Thus, a method that allows temporally-resolved tracking of spatial retrieval from LTM may provide a powerful tool for understanding human memory. Here, we present such a method, and show that it tracks both the accuracy and latency of memory-guided behavior. Moreover, we provide new evidence confirming a clear prediction of reinstatement models of LTM retrieval. The format of oscillatory activity during encoding into LTM is recapitulated during the subsequent retrieval of those memories.

## Acknowledgements

The authors would like to thank Brendan Colson, Nicholas Diaz, Jared Evans, Ariana Gale, Anubuv Gupta, Dylan Seitz, and William Ngiam for assistance with data collection, and Megan deBettencourt for helpful comments on the manuscript.

## References

Bohbot VD, Copara MS, Gotman J, Ekstrom AD. 2017. Low-frequency theta oscillations in the human hippocampus during real-world and virtual navigation. Nat Commun. 8:14415.

Bosch SE, Jehee JFM, Fernandez G, Doeller CF. 2014. Reinstatement of Associative Memories in Early Visual Cortex Is Signaled by the Hippocampus. J Neurosci. 34:7493–7500.

Brady TF, Konkle T, Gill J, Oliva A, Alvarez G a. 2013. Visual Long-Term Memory Has the Same Limit on Fidelity as Visual Working Memory. Psychol Sci. 24:981–990.

Brouwer GJ, Heeger DJ. 2009. Decoding and reconstructing color from responses in human visual cortex. J Neurosci. 29:13992–14003.

Cohen MX. 2014. Analyzing Neural Time Series Data: Theory and Practice, MIT Press.

Efron B, Tibshirani RJ. 1993. An introduction to the bootstrap. Chapman and Hall.

Ekstrom AD, Ranganath C. 2017. Space, time, and episodic memory: The hippocampus is all over the cognitive map. Hippocampus. 1–8.

Ester EF, Sprague TC, Serences JT. 2015. Parietal and Frontal Cortex Encode Stimulus-Specific Mnemonic Representations during Visual Working Memory. Neuron. 87:893–905.

Fan JE, Turk-Browne NB. 2013. Internal attention to features in visual short-term memory guides object learning. Cognition. 129:292–308.

Foster JJ, Bsales EM, Jaffe RJ, Awh E. 2017. Alpha-Band Activity Reveals Spontaneous Representations of Spatial Position in Visual Working Memory. Curr Biol. 1–8.

Foster JJ, Sutterer DW, Serences JT, Vogel EK, Awh E. 2016. The topography of alpha-band activity tracks the content of spatial working memory. 115:168–177.

Foster JJ, Sutterer DW, Serences JT, Vogel EK, Awh E. 2017. Alpha-Band Oscillations Enable Spatially and Temporally Resolved Tracking of Covert Spatial Attention. Psychol Sci. 28:929–941.

Harlow IM, Donaldson DI. 2013. Source accuracy data reveal the thresholded nature of human episodic memory. Psychon Bull Rev. 20:318–325.

Harlow IM, Yonelinas AP. 2014. Distinguishing between the success and precision of recollection. Memory. 8211:1–14.

Hindy NC, Ng FY, Turk-Browne NB. 2016. Linking pattern completion in the hippocampus to predictive coding in visual cortex. Nat Neurosci. 19:665–667.

Hsieh L, Ranganath C. 2014. Frontal midline theta oscillations during working memory maintenance and episodic encoding and retrieval. Neuroimage. 85:721–729.

Jafarpour A, Fuentemilla L, Horner AJ, Penny W, Duzel E. 2014. Replay of very early encoding representations during recollection. J Neurosci. 34:242–248.

Johnson JD, Price MH, Leiker EK. 2015. Episodic retrieval involves early and sustained effects of reactivating information from encoding. Neuroimage. 106:300–310.

Lins OG, Picton TW, Berg P, Scherg M. 1993. Ocular artifacts in recording EEGs and Event-Related potentials II: Source dipoles and source components. Brain Topogr. 6:65–78.

Maris E, Oostenveld R. 2007. Nonparametric statistical testing of EEG-and MEG-data. J Neurosci Methods. 164:177–190.

Morton NW, Kahana MJ, Rosenberg EA, Baltuch GH, Litt B, Sharan AD, Sperling MR, Polyn SM. 2013. Category-specific neural oscillations predict recall organization during memory search. Cereb Cortex. 23:2407–2422.

Morton NW, Polyn SM. 2017. Beta-band activity represents the recent past during episodic encoding. Neuroimage. 147:692–702.

Murray JG, Howie CA, Donaldson DI. 2015. The neural mechanism underlying recollection is sensitive to the quality of episodic memory: Event related potentials reveal a some-or-none threshold. Neuroimage. 120:298–308.

Nyhus E, Curran T. 2010. Functional role of gamma and theta oscillations in episodic memory. Neurosci Biobehav Rev. 34:1023–1035.

O’Keefe, Nadel L. 1978. The Hippocampus as a Cognitive Map, Oxford University Press.

Richter FR, Cooper RA, Bays PM, Simons JS. 2016. Distinct neural mechanisms underlie the success, precision, and vividness of episodic memory. Elife. 5:1–18.

Ritchey M, Wing EA, LaBar KS, Cabeza R. 2013. Neural Similarity Between Encoding and Retrieval is Related to Memory Via Hippocampal Interactions. Cereb Cortex. 23:2818–2828.

Saproo S, Serences JT. 2014. Attention Improves Transfer of Motion Information between V1 and MT. J Neurosci. 34:3586–3596.

Siegel M, Donner TH, Engel AK. 2012. Spectral fingerprints of large-scale neuronal interactions. Nat Rev Neurosci. 13:121–134.

Sprague TC, Ester EF, Serences JT. 2014. Reconstructions of Information in Visual Spatial Working Memory Degrade with Memory Load. Curr Biol. 24:2174–2180.

Sprague TC, Ester EF, Serences JT. 2016. Restoring Latent Visual Working Memory Representations in Human Cortex. Neuron. 91:694–707.

Sprague TC, Saproo S, Serences JT. 2015. Visual attention mitigates information loss in small- and large-scale neural codes. Trends Cogn Sci. 1–12.

Sprague TC, Serences JT. 2013. Attention modulates spatial priority maps in the human occipital, parietal and frontal cortices. Nat Neurosci. 16:1879–1887.

Stokes MG, Atherton K, Patai EZ, Nobre AC. 2012. Long-term memory prepares neural activity for perception. Proc…. 2011:188–197.

Suchow JW, Brady TF, Fougnie D, Alvarez G a. 2013. Modeling visual working memory with the MemToolbox. J Vis. 13:1–8.

Sutterer DW, Awh E. 2016. Retrieval practice enhances the accessibility but not the quality of memory. Psychon Bull Rev. 23:831–841.

Wagner AD, Shannon BJ, Kahn I, Buckner RL. 2005. Parietal lobe contributions to episodic memory retrieval. Trends Cogn Sci. 9:445–453.

Waldhauser GT, Braun V, Hanslmayr S. 2016. Episodic Memory Retrieval Functionally Relies on Very Rapid Reactivation of Sensory Information. J Neurosci. 36:251–260.

Waldhauser GT, Johansson M, Hanslmayr S. 2012. A/B Oscillations Indicate Inhibition of Interfering Visual Memories. J Neurosci. 32:1953–1961.

Watrous AJ, Ekstrom AD. 2014. The Spectro-Contextual Encoding and Retrieval Theory of Episodic Memory. Front Hum Neurosci. 8.

Watrous AJ, Fell J, Ekstrom AD, Axmacher N. 2015. More than spikes: Common oscillatory mechanisms for content specific neural representations during perception and memory. Curr Opin Neurobiol. 31:33–39.

Watrous AJ, Fried I, Ekstrom AD. 2011. Behavioral correlates of human hippocampal delta and theta oscillations during navigation. J Neurophysiol. 105:1747–1755.

Wilken P, Ma WJ. 2004. A detection theory account of change detection. J Vis. 4:1120–1135.

Wimber M, Maaß A, Staudigl T, Richardson-Klavehn A, Hanslmayr S. 2012. Rapid memory reactivation revealed by oscillatory entrainment. Curr Biol. 22:1482–1486.

Zhang W, Luck SJ. 2008. Discrete fixed-resolution representations in visual working memory. Nature. 453:233–235.

